# Modeling Risks and Mitigation Options for the Chronic Wasting Disease (CWD) in Scandinavia

**DOI:** 10.1101/2020.08.10.243782

**Authors:** Oskar Franklin, Elena Moltchanova, Andrey Krasovskiy, Florian Kraxner

## Abstract

Chronic wasting disease (CWD) is a contagious neural prion-disease affecting deer populations in North America with severe ecological and societal consequences. CWD is fatal and infectious prions spread and remain in the environment for many years even without animals present. The recent appearance of CWD in reindeers in Norway called for a drastic culling operation to prevent further spreading of the disease. This appears to have stopped the spreading of CWD among reindeers, but due to the persistence of CWD prions in the environment a reappearance of new cases among reindeer or other species in the future cannot be excluded. To evaluate the risks and the effectiveness of alternative management (monitoring and culling) options, we developed a model of CWD dynamics and management. The model includes stochastic population and spatial dynamics of the four relevant deer species in northern Sweden and Norway: reindeer (*Rangifer tarandus*), roe deer (*Capreolus capreolus*), red deer (*Cervus elaphus*) and moose (*Alces alces*). Transmission of CWD is modelled via direct contacts and via the environment. The model was parameterized and calibrated based on CWD studies from USA, data from the Norwegian CWD cases, and local deer population and vegetation data. The results indicate that without management, a CWD epidemic can be initiated by a single infected reindeer and would spread to other deer species. It would lead to dramatic population declines of reindeer and red deer and would also reduce the populations of roe deer and moose. The disease prevalence would stabilise at a about 50% after 50 years, as observed in some areas in the USA. A management strategy to cull only visibly sick animals, even with very efficient detection, cannot prevent a catastrophic development but merely slow the outbreak. To prevent an outbreak and the establishment of CWD it is necessary to cull all individuals, not only visibly sick ones, of an affected species in a relatively large area (30×30 km in our model) once a case is detected. Further, to prevent a slow buildup of CWD in the environment and eliminate the risk of outbreaks in the future it is necessary to expand this area of culling even further. Although the model has not yet been thoroughly validated due to scarcity of data, the results suggest that the drastic culling done in Norway was appropriate and necessary to prevent establishment of CWD and that further monitoring and potential culling is required to prevent outbreaks in the future.

## Introduction

The nature and history of chronic wasting disease (CWD) and its implications for the situation in Scandinavia is described in the comprehensive report by (Hansen et al. 2017). This knowledge was used to evaluate the options when CWD was first detected in Norway in 2016 and led to the controversial decision to cull an entire reindeer population in an attempt to exterminate the disease before it got established (Mysterud and Rolandsen 2018).

The history of spreading of CWD in North America shows that the disease initially spreads very slowly but steadily and after many years can reach a very high prevalence. In Colorado and Wyoming, models suggested 1% prevalence in 15-20 years, reaching 15% after 37-50 years (Miller et al. 2000) and reaching 40% or more in some areas, leading to deer population declines. Importantly, there are no known CWD outbreaks in which the pathogen has disappeared by itself after becoming established (Hansen et al. 2017). At the same time there are two cases (New York and Minnesota) of successful eradication of CWD under free-ranging conditions (open populations), based on massive, spatially targeted harvesting and the implementation of intense surveillance soon after the discovery of CWD. This suggests that an outbreak can only be stopped if proper measures are taken at an early stage of spreading.

CWD is transmitted both via direct contacts and environmental contamination. The relative importance of these transmission routes is not well known but it is reasonable to assume that direct transmission is more important in an early stage when environmental contamination is yet low (Almberg et al. 2011, Hansen et al. 2017). The rate of direct transmission depends on the number and frequency of contacts between individuals. Although the number of contacts increase with population density some studies suggest that this dependence is weak and therefore that CWD transmission is largely density independent (Habib et al. 2011), implying that population reduction is not an effective way to halt CWD spreading unless the whole population is eradicated (Hansen et al. 2017). However, we note that this conclusion may be premature because these studies did not account for environmental contamination and transmission, which will depend on population density.

In contrast to unselective population reduction, predation and selective hunting contribute to reducing the current proportion of CWD-infected individuals in the population. However, whether this is sufficient to reduce prevalence over time depends on the level of selectivity of predation and hunting and the rate of new CWD infections. In Illinois (USA) localized culling of deer in CWD infested areas stopped the increase in CWD prevalence whereas prevalence increased in Wisconsin where culling operations were discontinued (Manjerovic et al. 2014), suggesting that selective culling can be an effective mitigation measure.

The potential to stop or manage a CWD outbreak depends on the species involved. Although it is not yet confirmed, it is likely that CWD can infect all the Scandinavian deer species and potential efficiency of different management options may vary among species. The large and dynamic herds and social behavior of wild reindeer may lead to relative rapid transmission that cannot be managed by selective culling of smaller groups. Domestic reindeer are subject to high potential rates of transmission due to the frequent aggregation of animals, while on the other hand, it is possible to manage risks because they are largely under human control. Compared to reindeers, moose and roe deer are generally more solitary animals, which may limit a rise in prevalence of CWD and suggests that localized culling can be effective. However, if CWD infects roe deer in Norway, the high dispersal rate of yearling roe deer may rapidly spread the disease to new areas (Hansen et al. 2017), which suggests that restriction of movement, e.g. by fencing would be an important mitigation measure. An important measure to reduce CWD spreading in all species is the removal of hotspots - places where animals aggregate and transmit diseases at high rates, such as salt licks and feeding stations (Hansen et al. 2017, Western Association of Fish and Wildlife Agencies 2018).

In summary, management of CWD is a complex problem because multiple, highly uncertain processes and factors collectively determine the potential risk of an outbreak, the possibility to stop it, and which mitigation measures would be effective. In addition, most potential measures are costly and to have a chance to be effective, measures have to be taken at an early stage of disease spreading. In this situation, a conservative or trial and error approach is not an option. Instead a proactive approach based on predictive modeling of the potential disease dynamics and potential management options is required.

### Goals and research questions

The goal of the project is to develop a model with the capacity to evaluate the risks of CWD spreading and the potential effectiveness of mitigation measures. Based on the model we aim to evaluate the potential impact of management actions or no management in the short and the long term, for deer populations, for the spread and spatial distribution of CWD, and for our possibility to eradicate the disease. In the current model version, the management methods are monitoring and culling. Monitoring intensity influence the timing of disease detection and therefore the potential to eradicate it. We model different levels of culling, i.e. targeting only sick animals, culling all animals in an area affected, and culling animals also in areas around the affected area in order to eliminate potential undetected cases. Based on the model results, we also discuss whether the reindeer culling in Norway was an appropriate CWD management measure or whether less or even more drastic measured would have been preferable. However, it is important to keep in mind that this is an academic study and the degree to which the conclusions apply to the real case is yet uncertain due to the limited knowledge and scarcity of data regarding the epidemiology of CWD.

## Model

### Model overview

The model includes population dynamics of the relevant deer species: moose, roe deer, red deer, and reindeer, and the dynamics of CWD in the animal populations and in the environment (Fig. 1). The study area is divided into sub-areas (grid cells) and in each cell the environment is divided into different vegetation types (pine forest, spruce forest, deciduous forest, mixed forest, grassland, heath, and non-suitable habitat), which determines the spatial distribution of different species depending on their vegetation preferences (Table 2). In the current version of the model the vegetation distribution is identical in all cells. We model the spread of CWD between cells by letting the animals in one cell move and interact with animals and the environment in neighboring cells to an extent determined by their home range size relative to the cell size. In this model version we do not address the feedbacks between the deer populations and the vegetation dynamics as vegetation is assumed to be constant.

**Table 1.** Deer species parameters

| Parameter | Roe deer | Moose | Red deer | Reindeer |
| --- | --- | --- | --- | --- |
| Fertile ages (years) | 2 –8 | 2 -11<sup>3</sup> | 2-11<sup>2</sup> | 2 -11<sup>4</sup> |
| Maximum age (years) | 12<sup>1</sup> | 17<sup>3</sup> | 18<sup>2</sup> | 17<sup>4</sup> |
| Max number of calves | 3<sup>1</sup> | 2<sup>3</sup> | 1<sup>4</sup> | 1<sup>4</sup> |
| Group size | 1-4<sup>1</sup><br>(family group) | 1-3<sup>1</sup> (singels or<br>mother and<br>calves) | 5<sup>2</sup> | 100 (10 -200<sup>5</sup>) |
| Range size (km<sup>2</sup>, in<br>northern Sweden) | 5<sup>1</sup> | 50<sup>1</sup> | 50<sup>1</sup> | 300<sup>6</sup> |

References: <sup>1</sup> (Jarnemo et al. 2018), <sup>2</sup> British Deer Society, <sup>3</sup> (Kalen 2018), <sup>4</sup> Wikipedia, <sup>5</sup> Anna-Marja Kardik (pers.comm.), <sup>6</sup> Sameby Kall in Jämtland with a population of 2500 reindeers (Sametinget, https://www.sametinget.se/8816)

**Table 2.** Vegetation preferences of the deer species

| Vegetation | Roe deer | Moose | Red deer | Reindeer |
| --- | --- | --- | --- | --- |
| Broad-leaved forest | 0.32 | 0.26 | 0.31 | 0.16 |
| Mixed forest | 0.26 | 0.26 | 0.25 | 0.16 |
| Moors and heathland | 0.00 | 0.00 | 0.00 | 0.31 |
| Pastures+ natural grasslands | 0.32 | 0.16 | 0.25 | 0.31 |
| Pine forest | 0.06 | 0.26 | 0.13 | 0.06 |
| Spruce forest | 0.03 | 0.06 | 0.06 | 0.00 |

**Figure 1.**
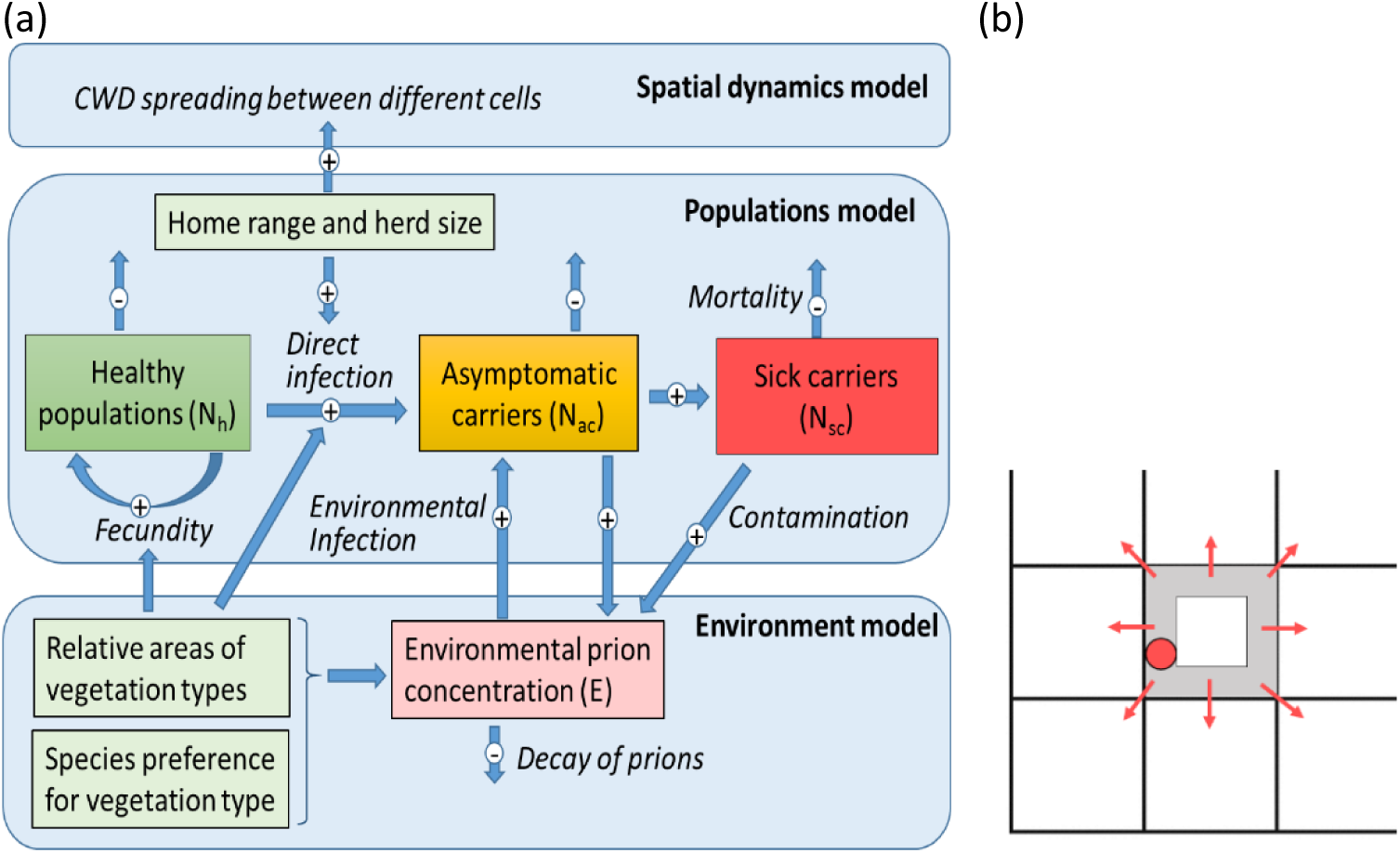
(**a**) Model structure. Variables (boxes), processes (italic text) and effects (blue arrows). (**b**) Spatial dynamics. The size of the home range of a species (red circle) relative to the cell size determines the fraction of its cell population (grey area) which can move back and forth or migrate out of the cell (red arrows).

The CWD dynamics is determined by infection, mortality of infected animals, environmental contamination, and prion decay in the environment. CWD can be transmitted via the environment or between individuals, for which the transmission rate depends on the number contacts, which in turn depends on the behavior of the animals in terms of home range size and herd size. Fecundity, mortality, and infection are stochastic processes, i.e. there is random variation. This means that the model is run multiple times to evaluate the probabilities of different outcomes.

### Population dynamics

The deer population dynamics is determined by rates of fecundity and mortality (Eq. 1a-c). Fecundity is density dependent so that the average number of calves declines with population density. Mortality related to natural causes (*m*_n_) is determined by the maximum life span. Additional mortality due to normal hunting (not CWD mitigation culling) and predation (*m*_h_) as well as the maximum population size (in the absence of mortality; *N*_max_) were estimated based on the current populations (Table 4) and the assumption that populations are at the levels of maximum sustainable yield, reflecting the underlying assumption that the populations currently are optimally managed. However, in the current application of the model the separation of normal hunting and natural mortality is not important, only their aggregated outcome matters and the assumption that current populations are in equilibrium.

**Table 3.** CWD parameters

| Parameter | Symbol | Values | References |
| --- | --- | --- | --- |
| CWD mortality | $m_{\text{cwd}}$ | 0.5 (95% CI: 0.34 – 0.61 yr<sup>-1</sup>) | (Miller et al. 2008) |
| Incubation time |  | 1.4 – 2 yr | (Zabel and Ortega 2017) |
| Transmission rate per contact (density dependent) | $\pi_{\text{cont}}$ | 0.00034 yr<sup>-1</sup><br>0.0328 yr<sup>-1</sup><br>0.0078 yr<sup>-1</sup> | (Wasserberg et al. 2009)<br>(Miller et al. 2006)<br>Estimate for reindeer* |
| Environmental infection rate | $p_{\text{env}}$ | 1.13 capita mass<sup>-1</sup> yr<sup>-1</sup> | (Miller et al. 2006) |
| Vertical transmission probability from cow to calf |  | 0.05 | (Gross and Miller 2001) |
| Environmental contamination | $c$ | 0.332 mass capita<sup>-1</sup> yr<sup>-1</sup> | (Miller et al. 2006) |
| Environmental prion decay | $d_E$ | 0.2 (0.1 – 5.76) yr<sup>-1</sup> | (Almberg et al. 2011).<br>(Miller et al. 2006) |

\*Based on CWD statistics in Norway: http://apps.vetinst.no/skrantesykestatistikk/NO/#omrade.

**Table 4.** Deer population densities. From (Jarnemo et al. 2018)

| Population density (km<sup>-2</sup>) | Roe deer | Moose | Red deer | Reindeer |
| --- | --- | --- | --- | --- |
| Area 1 (Western Jämtland)* | 0.8 | 0.4 | 0.4 | 0.83 |

\*see Fig. 2

Animals infected by CWD but not yet sick, asymptomatic carriers (*N*_ac_), are assumed to get sick with a constant probability (*p*_sc_) per year but otherwise not to differ from healthy individuals *N*_h_. Sick animals (*N*_sc_) have a high mortality (*m*_cwd_, Table 3).

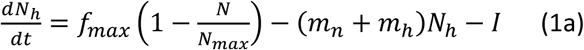

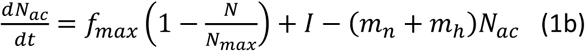

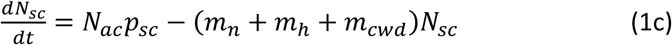

In the model stochastic versions of Eqs. 1a-c are used where both fecundity (first term in Eq 1) and mortalities (second term) are replaced by binomial distribution functions.

### Spatial movement and migration

We assume that the individuals in a cell can move to neighboring cells stochastically with a probability equal to the proportion of the population within a home range diameter of the border of the cell (Fig. 1). The probability of moving horizontally or vertically is twice as high as the probability of moving diagonally. Half of the moving individuals will stay in their new cell and the rest will merely share their time spent between their current and their neighboring cells. This leads to mingling among populations in different cells, which contribute to the transmission of CWD. The number and sizes of grid cells can be varied in the model and in the reported simulations we used a 10 x 10 cells grid with cell size 30 x 30 km.

### CWD transmission

#### Direct transmission between individuals

The transmission of the disease can occur from four sources: (i) from within the herd, (ii) from animals outside the herd but within the same cell, (iii) from the animals in the neighbouring cells, and (iv) from the environment (prion contamination). We assume that the individual-to-individual transmission can only occur within a species, whereas all the species contribute to the environmental prion load.

We consider an individual cell with a population of *N* individuals of a particular species, divided into *h* herds of size *s*, and assume *N*_*ac*_ *+N*_*sc*_ out of *N* are carriers. Furthermore, *A*_*r*_ denotes the home range size of the herd. Then the expected number of contacts for a focal individual from within its herd is *s*-1. The expected number of contacts from within the cell but outside the herd is the number of individuals in other herds whose ranges intersect with the focal individual, i.e.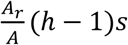, where *A*_c_ is the area of the cell. The infected individuals are assumed to be spread uniformly throughout the population (somewhat inaccurate, given the herding), so the number of expected contacts with infected animals are

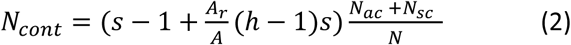

The number of contacts should be interpreted as the expected number of different individuals that a focal individual comes into contact with during one year. The probability of contracting the disease from these contacts becomes

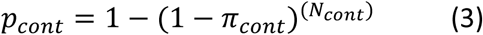

where *π*_cont_ is the probability of transmission via each individual-to-individual contact.

#### Environmental contamination and transmission

We assume that each infected animal contaminate the environment at a constant rate (*c*). The contamination (*E*, eq. 4) is distributed among vegetation types (index *v*) in proportion to the vegetation preferences of each species (*pr*_v_, Table 2). This implies that more preferred vegetation types will be more contaminated and that inter species transmission of CWD is more likely between species that share vegetation preferences. Prions in the environment are assumed to decay at a constant rate (*d*_E_).

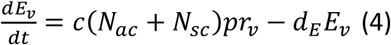

The probability of environment-to-individual transmission is a monotonic increasing function of the environmental load (*E*), taking values between 0 and 1:

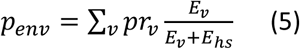

#### Infection rate

The total probability of getting CWD is equal to 1 minus the probability of not getting it either from contacts or the environment:

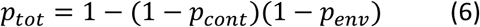

The number of new cases of CWD per time (*I*) will then follow a binomial distribution:

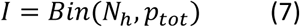

### CWD Management

Two aspects of management are currently included in the model, monitoring and culling. Monitoring effort is modelled with the parameter *θ*, which is the probability of detecting an infected animal (a case). We assume that only the symptomatic carriers can be detected. The culling currently includes four options:

0. No culling
1. Cull the sick animals: If a sick animal is detected it is culled.
2. Cull the cell: If a sick animal is detected, then all the animals of the same species in the cell are culled.
3. Cull the cell and the neighbourhood: If a sick animal is detected, then all the animals of the same species are culled in the cell and in its neighbourhood.

The neighborhood is defined as all cells around a cell (Fig. 1b).

### Data and model parameters

Many parameters and population information for all species except reindeers were taken from the report Hjortvilt i Sverige (Jarnemo et al. 2018) but also from various other sources (Tables 1 -4). CWD parameters were taken from literature, mainly based on American studies. The contact infection probability (*π*_cont_) was observed to be very important for the simulation results and at the same time to strongly vary between studies from 0.00034 (Wasserberg et al. 2009) to 0.0328 (Miller et al. 2006). Thus, instead of using these values we estimated *π*_cont_ based on the observed populations and CWD cases in Norwegian reindeers available at http://apps.vetinst.no/skrantesykestatistikk/NO/#omrade. In this estimation we assumed that the average group size of reindeers was 100 animals and we tested two alternatives for the number of contacts among groups (or home range overlap), A larger group size or larger home range leads to lower *π*_cont_ and vice versa but both estimates lie between the values reported in the American studies. Because the results were qualitatively similar for the two values of *π*_cont_ we present results only for one value of *π*_cont_ = 0.0078 below.

### Model implementation and interface

The current version of the model is implemented in MathCad (version 13; https://en.wikipedia.org/wiki/Mathcad#Overview). All different culling scenarios (See section CWD Management) are evaluated and the outputs are saved in a data file, which is used by a user interface to display results for selected scenarios and parameters. This approach makes it possible to quickly compare different alternatives without having to wait for the model complete a new simulation for each setting one at a time.

We evaluated the model for an area in Sweden near the Norwegian border (Fig. 2), initialized with data on the populations (Table 4) and the vegetation (Table 5). This area was chosen because of its proximity to Norway (where CWD has been detected) and because it contains all the most relevant deer species (reindeer, moose, red deer, and roe deer).

**Table 5.** Vegetation types in the studied area ^#^

| Vegetation type | share |
| --- | --- |
| Broad-leaved forest | 5.3% |
| Mixed forest | 13.8% |
| Moors and heathland | 21.1% |
| Pastures+ natural grasslands | 1.3% |
| Pine forest* | 2.9% |
| Spruce forest* | 27.3% |
| The rest (everything with no food for deer) | 28.4% |

\*Data taken from (SLU 2018) and https://land.copernicus.eu/pan-european.
<sup>#</sup>see Fig. 2

**Figure 2.**
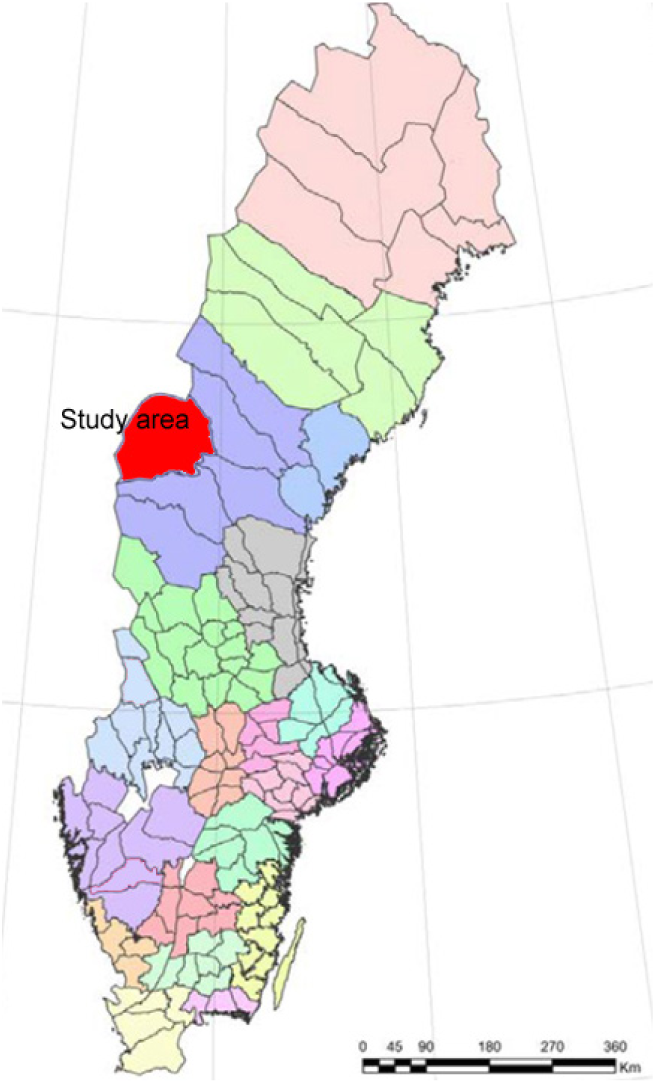
Map of Sweden with the area studied (western Jämtland) marked in red.

Each simulation was started with one sick reindeer entering a corner of the study area (as yet only reindeers have been confirmed to carry the contagious version of CWD in Norway). For each parameter setting 100 iterations were simulated and the mean values for each time point, 95% confidence intervals, and probability of 0 infected animals (elimination of CWD), is evaluated for each cell and each species and over all cells, for each year simulated.

## Results

We first evaluated the scenario 0 with no management (Fig. 3), serving as a baseline to compare with the management alternatives. Without management the model predicts an exponential increase in the number of CWD cases and the spatial area affected. Initially, mainly reindeer are infected while later the disease is spread to the other species via the environment. The rapid spread of CWD among reindeer results in a steep population decline of reindeers and subsequently also of the other species. This population decline reduces the buildup of prions in the environment and the number of new infections. This eventually leads to a partial recovery of moose and roe deer populations, which stabilize at a level below 50% of initial populations and with a CWD prevalence of ca 50%. In contrast, the reindeer and red deer populations do not recover and their populations remain at only a few hundred over the entire study area.

**Figure 3.**
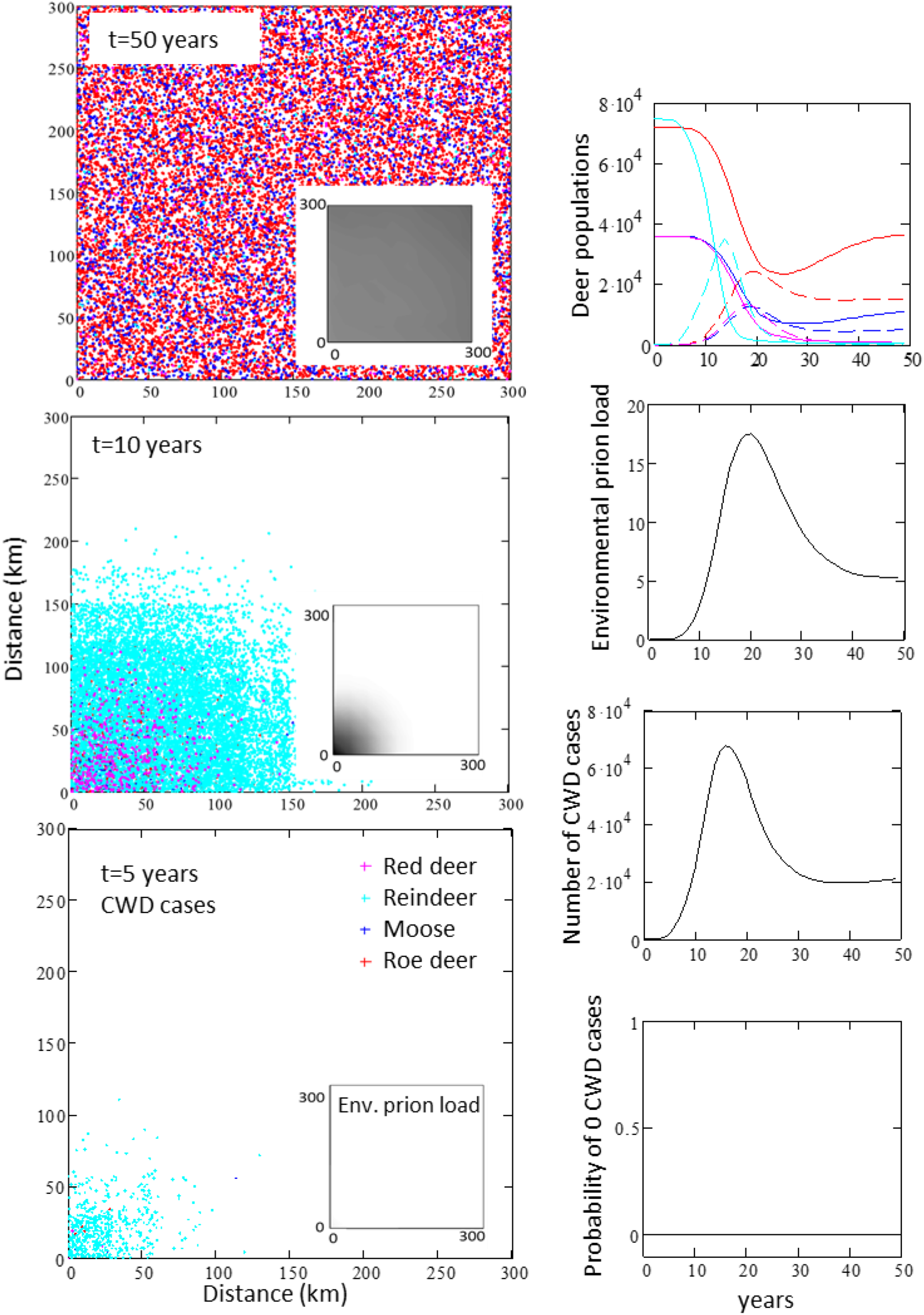
Baseline scenario with no management. Left panels: Spatial distribution of CWD cases. Inserts in left panels: Spatial distribution of environmental CWD load (shading). Right panels from top: Temporal development of deer populations, environmental prion load, number of CWD cases, and the probability of CWD elimination (no CWD cases). All values (except probability of 0 CWD cases) are means values of 100 simulations.

In management scenario 1 all detected sick animals were culled with a detection probability of 90% (Fig. 4). This management option cannot prevent a CWD outbreak but reduces the number of cases and the long term CWD prevalence with about 50% compared to no management. Red deer and reindeer are able to persist at population levels of about 3000 (5 −10% of pre CWD populations).

**Figure 4.**
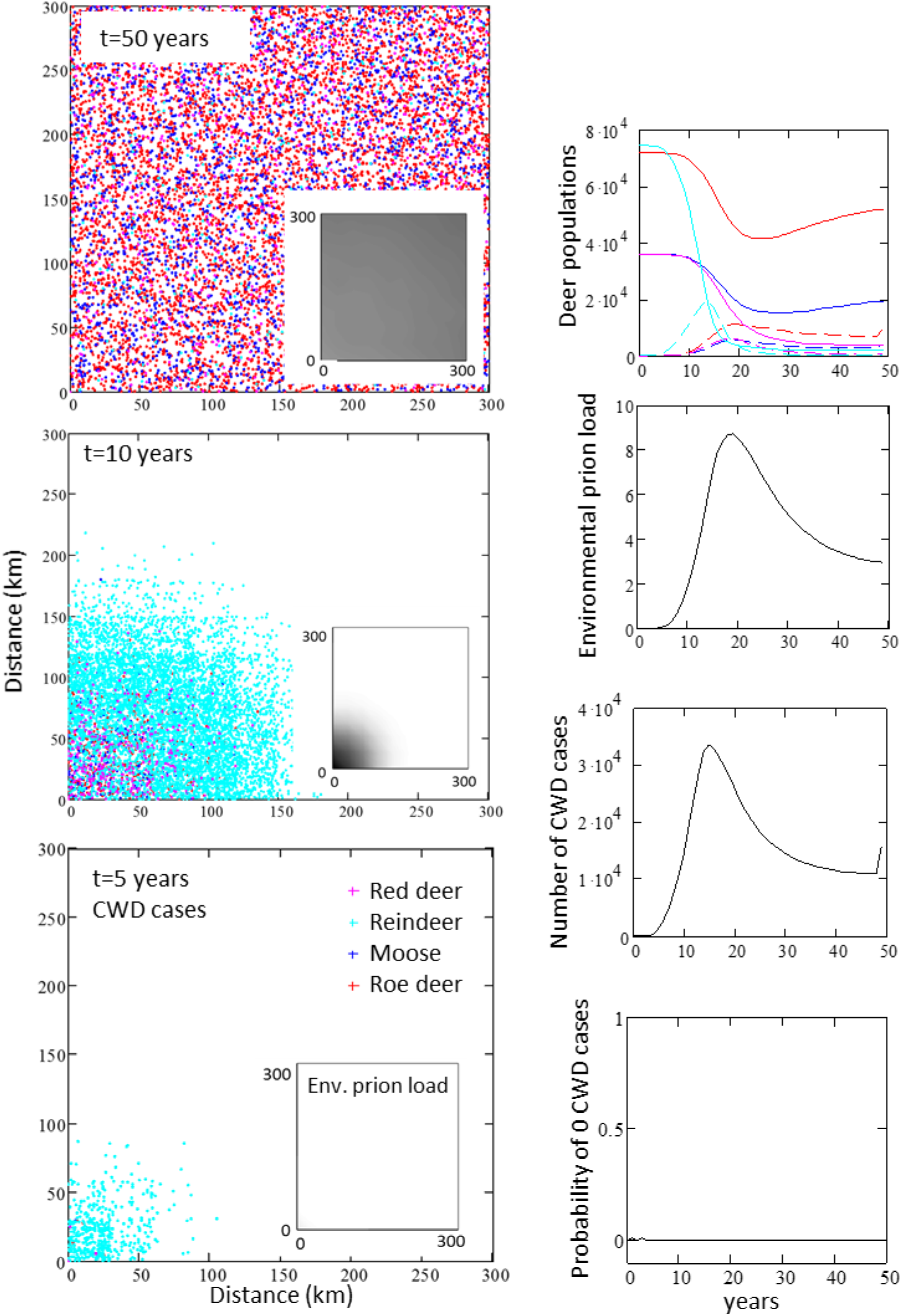
Management scenario 1 – culling only infected animals, detection probability 90%. Left panels: Spatial distribution of CWD cases. Inserts in left panels: Spatial distribution of environmental CWD contamination (shading). Right panels from top: Temporal development of deer populations, environmental prion load, CWD cases, and the probability of CWD elimination (no CWD cases). All values (except probability of 0 CWD cases) are means values of 100 simulations.

In management scenario 2 all animals of a detected infected species within the cell are culled, i.e. also healthy and infected animals that are not yet sick are culled (Fig. 5). This locally drastic culling only marginally reduces the total populations in the study area but has a dramatic effect on the CWD epidemic. The maximum number of CWD cases in one year is reduced from 10s of thousands to only a few sporadic cases, and there is a 50% probability that no cases are present in any given year. However, the prion contamination in the environment is slowly increasing and spreading across the area while at the same time the number of CWD cases is stable and even increasing in the last years of the simulation. This suggests that CWD will persist at low levels and may increase in the future.

**Figure 5.**
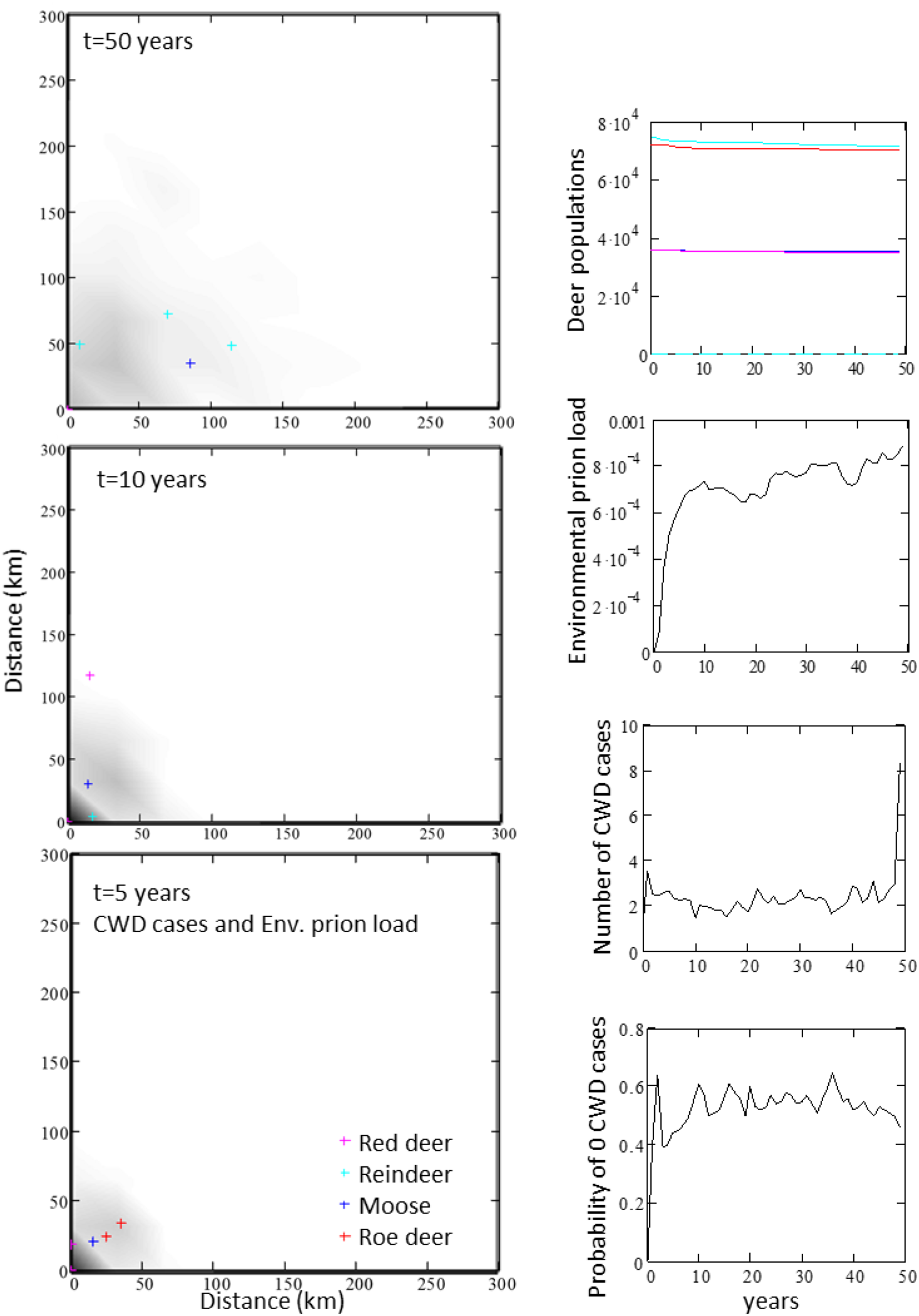
Management scenario 2 – culling all animals of an infected species in the cell, detection probability 90%. Left panels: Spatial distribution of CWD cases on top of environmental prion load (shading). Right panels from top: Temporal development of deer populations, environmental prion load, CWD cases, and the probability of CWD elimination (no CWD cases). All values (except probability of 0 CWD cases) are means values of 100 simulations.

Management scenario 3 is identical to scenario 2 except that additionally all individuals of a detected CWD affected species in neighboring cells are culled, i.e. deer of an affected species are culled over an up to 9 times larger area than in scenario 2 (Fig. 6). This management effectively mitigates an outbreak and spatial spreading of CWD. Only in the first few years new CWD cases are likely to appear while the probability of total CWD elimination increases over time to about 90% after 20 years and 95% after 35 years. The concurrent decline of environmental prion load suggests that CWD will eventually be completely eradicated under this scenario.

**Figure 6.**
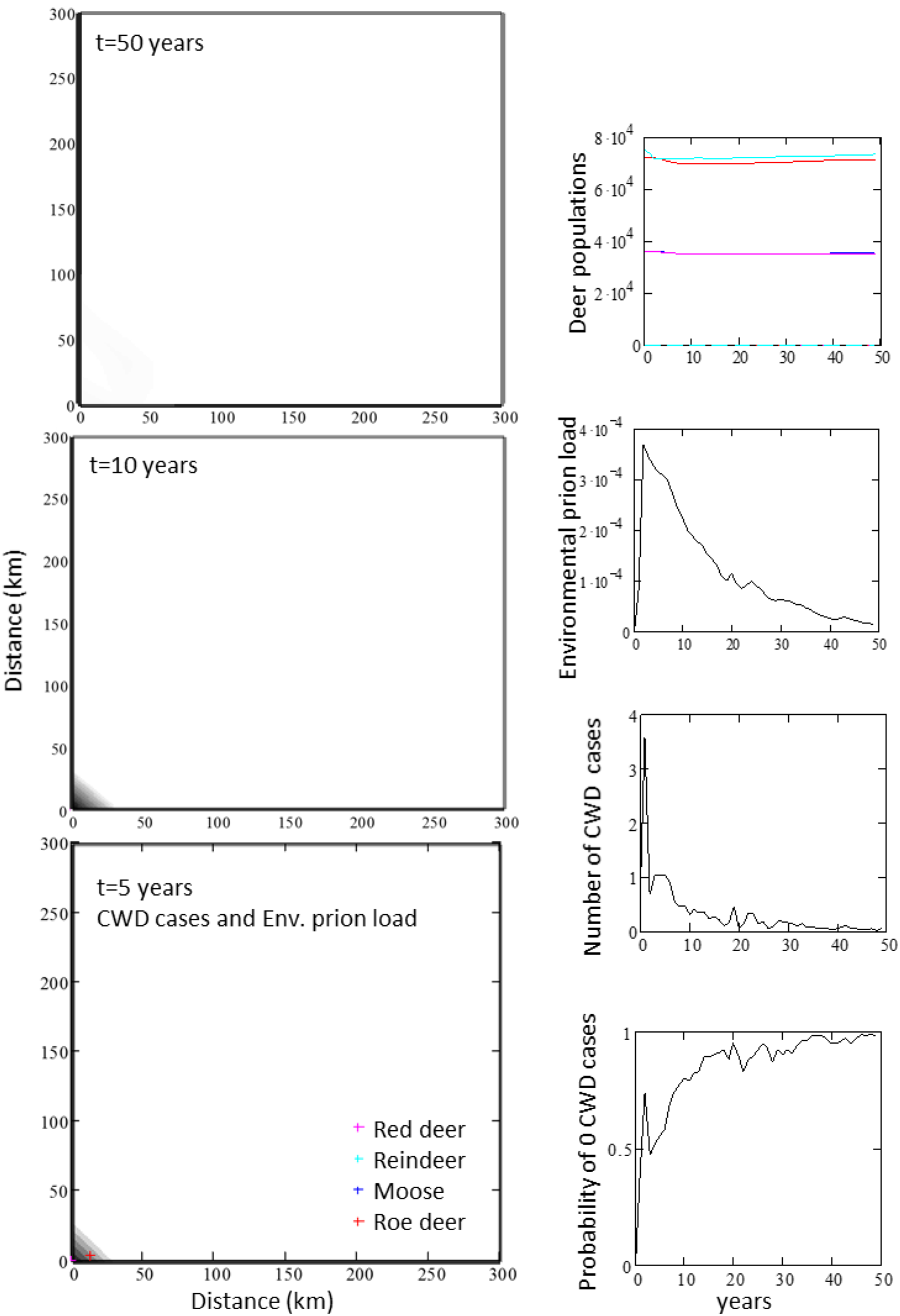
Management scenario 3 – culling all animals of an infected species in the cell and it neighboring cells, detection probability 90%. Left panels: Spatial distribution of CWD cases on top of environmental prion load (shading). Right panels from top: Temporal development of deer populations, environmental prion load, CWD cases, and the probability of CWD elimination (no CWD cases). All values (except probability of 0 CWD cases) are means values of 100 simulations.

## Discussion and conclusions

The model simulations suggest that a single infected reindeer may cause a CWD outbreak if management actions are not taken. Over time this scenario would lead to a persistent high prevalence of CWD and strongly reduced populations of all the deer species, and especially reindeer and red deer. Reindeer and red deer may be more sensitive to CWD than moose and red deer because of their larger herd sizes and their lower maximum fecundity (Table 1). However, the results should be interpreted with care because of the limited knowledge available on CWD and the resulting uncertainty in some important parameters, such as the probabilities of transmission of CWD by contact and from the environment, and the rate of decay of prions in the environment. The probability of an outbreak and its rate of increase is particularly sensitive to the contact transmission probability (*π*_cont_). To obtain the most relevant estimate possible for contact transmission probability we used the data from the reindeer in CWD affected area in Norway, http://apps.vetinst.no/skrantesykestatistikk/NO/#omrade). In addition to the results shown for *π*_cont_ = 0.0078 we tested the effect of assuming more contacts between herds, which led to a value of *π*_cont_ = 0.0011. The effect of the seven-fold lower value was a significantly slower progression of the outbreak under scenario 0 and 1 whereas the long-term outcomes and the effect of management options 2 and 3 did not change qualitatively. Although additional sensitivity analysis and testing against observed data remains to be done, the validity of the model is supported at least to some extent by the predicted long term prevalence and population decline, which is comparable to observations in some areas in USA where CWD has been present since a long time (DeVivo et al. 2017).

The evaluation of management options clearly shows that a strategy of culling only visibly sick animals is not an effective way to prevent an outbreak, even if sick animals are effectively detected. This is not surprising because of the CWD incubation time, which allows animals to spread the disease before they get sick and can be detected. Therefore, it is also not surprising that management scenario 2 where all individuals of an affected species in a cell (30×30 km) are culled after a case is detected, is much more effective. The model simulations indicate that with this management only sporadic cases of CWD will occur although CWD will persist over time (Fig. 5). A somewhat similar management strategy with comparable results has been applied in USA where deer were culled in CWD affected areas of similar size to our cells (ca 1000 km^2^) (Manjerovic et al. 2014). If this culling strategy (scenario 2) is extended also to surrounding cells (scenario 3), the model predicts that CWD will be eliminated with a probability of 95% after 35 years (Fig. 6). To our knowledge, such an extensive culling regime has yet never been applied in practice.

Regarding the CWD situation in Norway, the model results suggests that that the management strategy applied was appropriate and is likely to have prevented, at least temporarily, the initiation of a larger scale CWD outbreak and its far reaching consequences (Mysterud and Rolandsen 2018). The results also suggest that further monitoring is essential and that similar culling actions are required if new cases are detected in order to prevent a CWD outbreak in the future. Based on the model results, we propose that to reduce uncertainty about the risk of CWD outbreaks, further research is needed on how CWD transmission between animals depends on behavior (e.g. herd size and home range) and species. For more robust estimates of the long-term risks of CWD more knowledge is needed on the accumulation, infectiousness, and persistence of CWD prions in the environment. Further modeling studies could be used to evaluate additional management options, such as fencing (Mysterud and Rolandsen 2019), and how landscape and migration patterns influence the risks of CWD. In a societal perspective, a reliable model is essential to able to compare among alternative management options in term of effectiveness versus costs, as well as social and ecological impacts.

Although additional testing and evaluation based on observations and by other researchers and experts are needed to further validate the model and its results, we believe that the study demonstrates that this modeling approach is useful for evaluating the risks of CWD and the possible management options. Further, the model can be extended to integrate additional aspects of wildlife and society, such as other diseases, forestry, hunting, predators, and traffic. This would facilitate holistic analyses of wildlife and ecosystem management which may help to provide a more common understanding among different stakeholders.

## Notes

### Competing Interest Statement

The authors have declared no competing interest.

